# Crystallographic and electrophilic fragment screening of the SARS-CoV-2 main protease

**DOI:** 10.1101/2020.05.27.118117

**Authors:** Alice Douangamath, Daren Fearon, Paul Gehrtz, Tobias Krojer, Petra Lukacik, C. David Owen, Efrat Resnick, Claire Strain-Damerell, Anthony Aimon, Péter Ábrányi-Balogh, José Brandaõ-Neto, Anna Carbery, Gemma Davison, Alexandre Dias, Thomas D Downes, Louise Dunnett, Michael Fairhead, James D. Firth, S. Paul Jones, Aaron Keely, György M. Keserü, Hanna F Klein, Mathew P. Martin, Martin E. M. Noble, Peter O’Brien, Ailsa Powell, Rambabu Reddi, Rachael Skyner, Matthew Snee, Michael J. Waring, Conor Wild, Nir London, Frank von Delft, Martin A. Walsh

## Abstract

COVID-19, caused by SARS-CoV-2, lacks effective therapeutics. Additionally, no antiviral drugs or vaccines were developed against the closely related coronavirus, SARS-CoV-1 or MERS-CoV, despite previous zoonotic outbreaks. To identify starting points for such therapeutics, we performed a large-scale screen of electrophile and non-covalent fragments through a combined mass spectrometry and X-ray approach against the SARS-CoV-2 main protease, one of two cysteine viral proteases essential for viral replication. Our crystallographic screen identified 71 hits that span the entire active site, as well as 3 hits at the dimer interface. These structures reveal routes to rapidly develop more potent inhibitors through merging of covalent and non-covalent fragment hits; one series of low-reactivity, tractable covalent fragments was progressed to discover improved binders. These combined hits offer unprecedented structural and reactivity information for on-going structure-based drug design against SARS-CoV-2 main protease.

## Introduction

A novel coronavirus, SARS-CoV-2, the causative agent of COVID-19 (Wu et al., 2020, Kucharski et al., 2020, Zhu et al., 2020), has resulted in over one million confirmed cases and in excess of 300,000 deaths across 188 countries as of mid-May 2020 (Dong et al., 2020). SARS-CoV-2 is the third zoonotic coronavirus outbreak after the emergence of SARS-CoV-1 in 2002 and the Middle East Respiratory Syndrome (MERS-CoV) in 2012 (Bermingham et al., 2012, Kuiken et al., 2003, Zaki et al., 2012). SARS-CoV-2 is a large enveloped, positive sense, single stranded RNA Betacoronavirus. The viral RNA encodes two open reading frames that, through ribosome frame-shifting, generates two polyproteins pp1a and pp1ab (Bredenbeek et al., 1990). These polyproteins produce most of the proteins of the replicase-transcriptase complex (Thiel et al., 2003). The polyproteins are processed by two viral cysteine proteases: a Papain-like protease (PL^pro^) which cleaves three sites, releasing non-structural proteins nsp1-3 and a 3C-like protease, also referred to as the main protease (M^pro^), that cleaves at 11 sites to release non-structural proteins (nsp4-16). These non-structural proteins form the replicase complex responsible for replication and transcription of the viral genome and have led to M^pro^ and PL^Pro^ being the primary targets for antiviral drug development (Hilgenfeld, 2014).

Structural studies have played a key role in drug development and were quickly applied during the first coronavirus outbreak. Early work by the Hilgenfeld group facilitated targeting the M^pro^ of coronarviruses (Hilgenfeld, 2014), and this was accelerated during the 2002 SARS-CoV-1 outbreak, leading to crystal structures of SARS-CoV-1 M^pro^ and inhibitor complexes (Ghosh et al., 2007, Verschueren et al., 2008, Yang et al., 2005, Yang et al., 2003). Coronavirus M^pro^ active sites are well conserved (Anand et al., 2003, Hegyi and Ziebuhr, 2002, Stadler et al., 2003, Xue et al., 2008, Yang et al., 2005, Zhang et al., 2020b) and those of enteroviruses (3C^pro^) are functionally similar, which has led to efforts to develop broad spectrum antivirals. The most successful have been peptidomimetic α-ketoamide inhibitors (Zhang et al., 2020a), which have been used to derive a potent α-ketoamide inhibitor that may lead to a successful antiviral (Zhang et al., 2020b).

To date, no drugs targeting SARS-CoV-2 have been verified by clinical trials and treatments are limited to those targeting disease symptoms. To contribute to future therapeutic possibilities, we approached the SARS-CoV-2 M^pro^ as a target for high throughput drug discovery using a fragment-based approach (Thomas et al., 2019). We screened against over 1250 unique fragments leading to the identification of 74 high value fragment hits, including 23 non-covalent and 48 covalent hits in the active site, and 3 hits at the vital dimerization interface. Here, these data are detailed along with potential ways forward for rapid follow-up design of improved, more potent, compounds.

## Results

### M^pro^ crystallizes in a ligand-free form that diffracts to near-atomic resolution

We report the apo structure of SARS-CoV-2 M^pro^ with data to 1.25 Å. The construct we crystallised has native residues at both N- and C--terminals, without cloning truncations or appendages which could otherwise interfere with fragment binding. Electron density is present for all residues, including 26 alternate conformations, many of which were absent in previous lower resolution crystal structures. The protein crystallised with a single protein polypeptide in the asymmetric unit, and the catalytic dimer provided by a symmetry-related molecule. The structure aligns closely with the M^pro^ structures from SARS-CoV-1 and MERS (rmsd of 0.52 Å and 0.97 Å respectively). The active site is sandwiched between two β-barrel domains, I (residue 10-99) and II (residue 100-182) (Figure 1A). Domain III (residue 198-306), forms a bundle of alpha helices and is proposed to regulate dimerization (Shi and Song, 2006). The C-terminal residues, Cys300-Gln306, wrap against Domain II. However, the C terminal displays a degree of flexibility and wraps around domain III in the N3 inhibitor complex (Shi and Song, 2006) (PDB ID 6LU7). His41 and Cys145 comprise the catalytic dyad and dimerisation completes the active site by bringing Ser1 of the second dimer protomer into proximity with Glu166 (Figure 1B). This aids formation of the substrate specificity pocket and the oxyanion hole (Hilgenfeld, 2014). Subsites have previously been identified in the active site based on interactions with peptide-based inhibitors and are shown in figure 1B (Jin et al., 2020, Zhang et al., 2020b). Comparisons with peptide-based inhibitor complexes (Jin et al., 2020, Zhang et al., 2020b) suggest a degree of active site plasticity. In particular, the C-alphas of Met49, Pro168, Gln189 respectively show movements of 2.8 Å, 1.4 Å, and 1.2 Å in comparison to the α-ketoamide inhibitor bound M^pro^ structure (Zhang et al., 2020b) (PDB ID 6y2f, Figure 1B).

**Figure 1.**
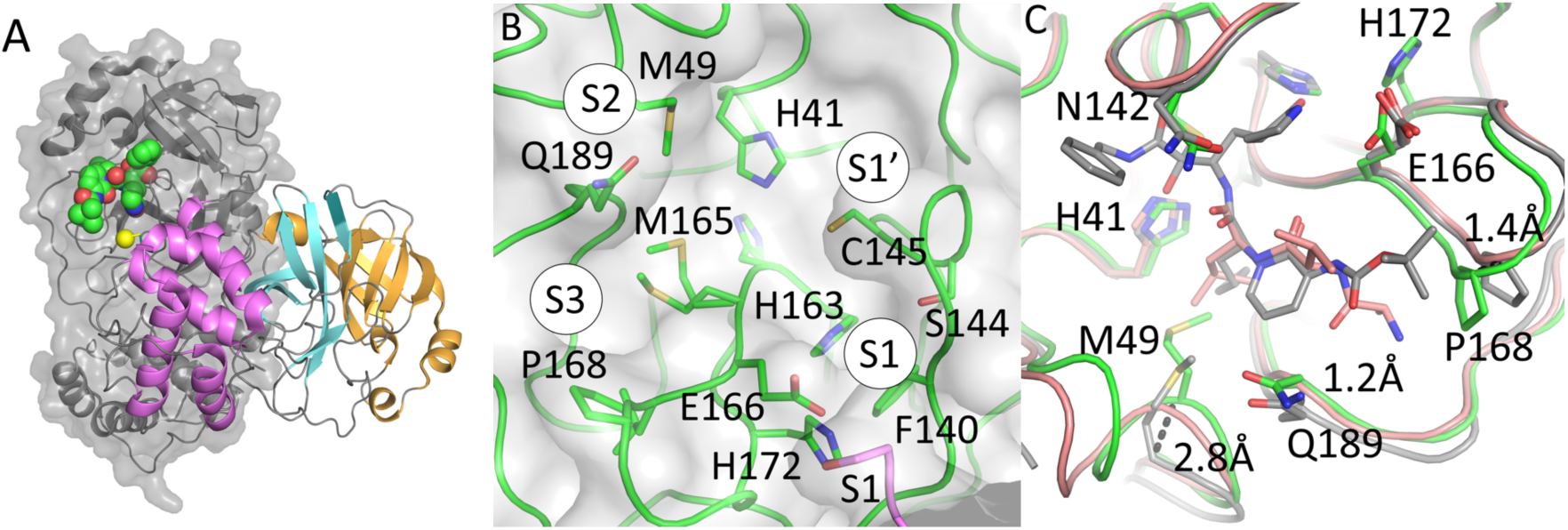
The crystal structure of ligand free M^pro^ is amenable to X-ray fragment screening. **A.** cartoon representation of the M^pro^ dimer. The nearmost monomer is shown with secondary structure features coloured to demarcate domains I, II, and III, in orange, cyan, and violet respectively. The active site of the rear monomer is indicated by the presence of a peptide-based inhibitor in green generated by aligning the ligand-free structure with pdb 6Y2F (Zhang et al., 2020b). A yellow sphere indicates Ser1 from the dimer partner that completes the active site. **B.** Residues of the active site are labelled, and subsites involved in ligand binding are shown with circles. **C.** Active site plasticity is observed when comparing apo structure to peptide inhibitor bound structures (green – Apo, grey – 6Y2F, pink 6LU7). Displacement distances associated with loop movements are indicated.

The crystal form is well-suited for crystallographic fragment screening: although the percentage of solvent is very low for a protein crystal, approximately 20%, nevertheless clear channels are present that allow access to the active site through diffusion. Moreover, the tight packing and strong innate diffraction mean crystals are resistant to lattice disruption and degradation of diffraction by DMSO solvent when adding solubilised fragments to the crystallization drop.

### Combined MS and crystallographic fragment screens reveal new binders of M^pro^

Cysteine proteases are attractive targets for covalent inhibitors. To identify covalent starting points, we screened our previously described library of ∼1000 mild electrophilic fragments (Resnick et al., 2019) against M^pro^ using intact protein mass spectrometry. Standard conditions of 200 µM per electrophile for 24 hours at 4 °C did not allow discrimination between hits. Screening at more stringent conditions (5 µM per electrophile; 1.5 hours; 25°C) resulted in 8.5% of the library labelling above 30% of protein (Table S1a). These hits revealed common motifs, and we focused on compounds which offer promising starting points.

Compounds containing *N*-chloroacetyl-*N*’-sulfonamido-piperazine or *N-*-chloroacetylaniline motifs were frequent hitters. Such compounds can be highly reactive. Therefore, we chose series members with relatively low reactivity for follow up crystallization attempts. For another series of hit compounds, containing a *N*-chloroacetyl piperidinyl-4-carboxamide motif (Table S2) which displays lower reactivity and were not frequent hitters in previous screens, we attempted crystallization despite their absence of labelling in the stringent conditions.

While mild electrophilic fragments are ideal for probing the binding properties around the active site cysteine, their small size prevents extensive exploration of the substrate binding pocket. We performed an additional crystallographic fragment screen to exhaustively probe the M^pro^ active site, and to find opportunities for fragment merging or growing. The 68 electrophile fragment hits were added to crystals along with a total of 1176 unique fragments from 7 libraries (Table S3). Non-covalent fragments were soaked (Collins et al., 2017), whereas electrophile fragments were both soaked and co-crystallized as previously described (Resnick et al., 2019), to ensure that as many of the mass spectrometry hits as possible were structurally observed. A total of 1742 soaking and 1139 co-crystallization experiments resulted in 1877 mounted crystals. While some fragments either destroyed the crystals or their diffraction, 1638 datasets with a resolution better than 2.8 Å were collected. The best crystals diffracted to better than 1.4 Å, but diffraction to 1.8 Å was more typical, and no datasets worse than 2.8 Å were included in analysis (Figure S2). We identified 96 fragment hits using the PanDDA method (Pearce et al., 2017), all of which were deposited in the Protein Data Bank (Table S2), but also immediately released through the Diamond Light Source website (https://www.diamond.ac.uk/covid-19.html), along with all protocols and experimental details. A timeline of experiments is shown in figure 2.

**Figure 2.**
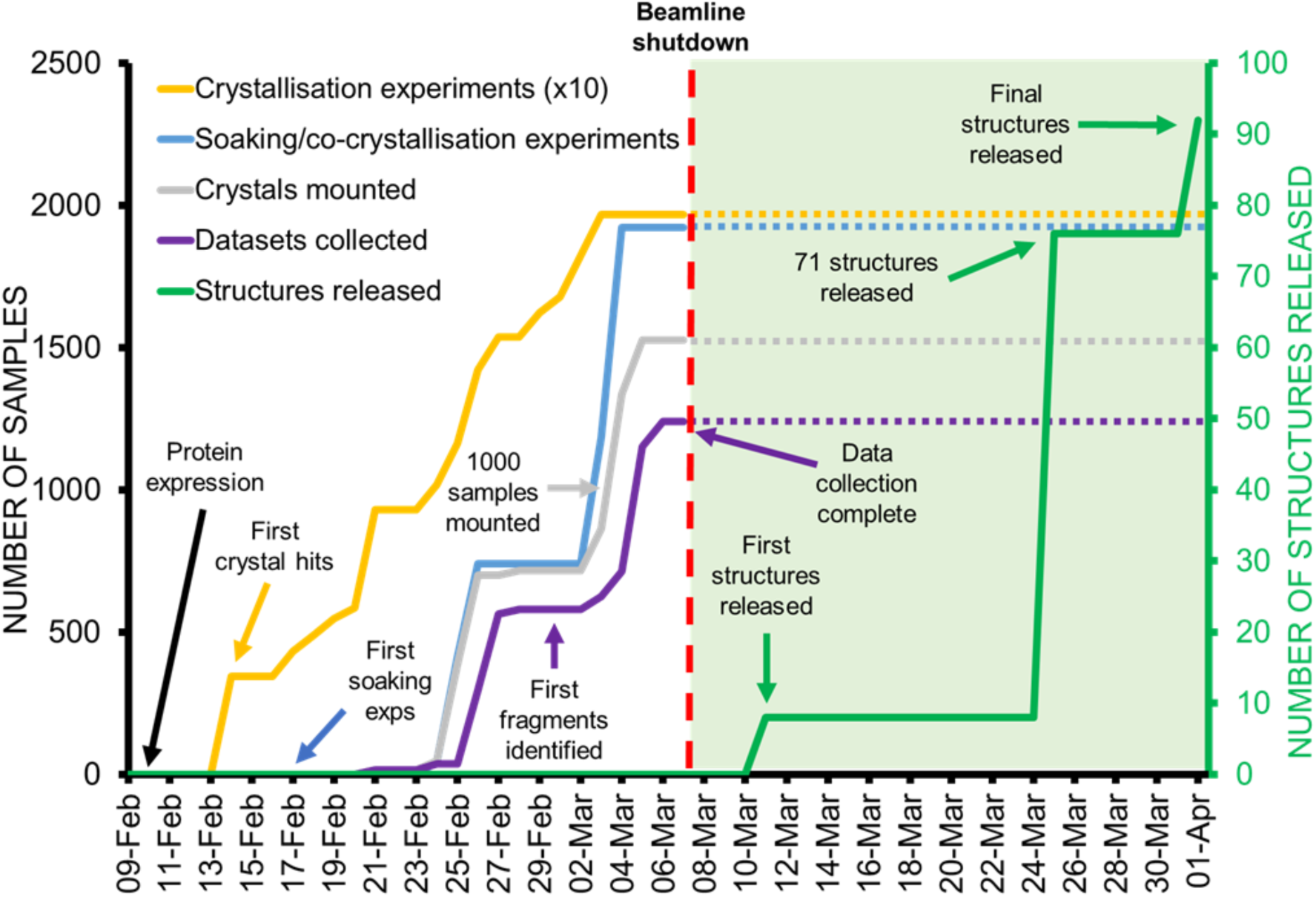
Timeline of crystallographic fragment screen.

### Non-covalent fragment hits reveal multiple targetable sub-sites in the active site

This unusually large screen identified 23 structurally diverse fragments that bind non-covalently and extensively sample features of the M^pro^ active site and its specificity pockets/subsites (figure 1), along with 3 hits exploring the dimer interface.

#### Active-site fragments

Eight fragments were identified that bind in the S1 subsite and frequently form interactions with the side chains of the key residues His163, through a pyridine ring or similar nitrogen containing heterocycle, and Glu166 through a carbonyl group in an amide or urea moiety (Figure 3). Several also reach across into the S2 subsite.

**Figure 3.**
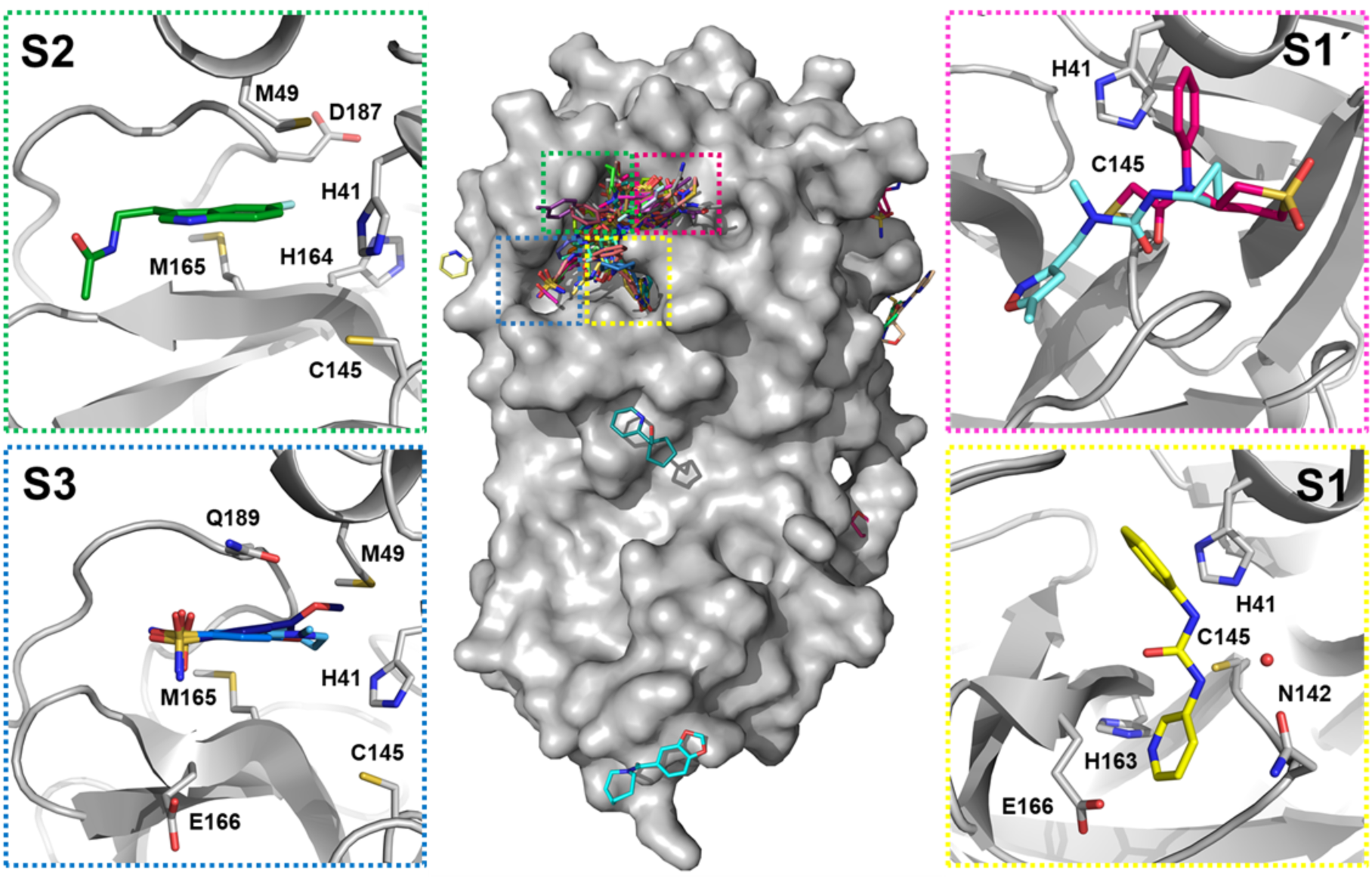
Bound fragments sample the active site comprehensively. The central surface representation is of the M^pro^ monomer with all fragment hits shown as sticks, and active site subsites highlighted by coloured boxes. Each subsite expanded along with a selection of hits to demonstrate common features and interactions. **S1**: Z44592329 (x0434); **S1’**: Z369936976 (x0397) in aquamarine and PCM-0102372 (x1311) in magenta bound to active site cysteine; **S2**: Z1220452176 (x0104); **S3**: Overlay of Z18197050 (x0161), Z1367324110 (x0195) and NCL-00023830 (x0946).

Subsite S2 has previously demonstrated greater flexibility in comparison to the other subsites, adapting to smaller substituents in peptide-based inhibitors but with a preference for leucine or other hydrophobic residues (Zhang et al., 2020b). Many fragments bound at this location, which we termed the “aromatic wheel” because of a consistent motif of an aromatic ring forming hydrophobic interactions with Met49 or π-π stacking with His41, with groups variously placed in 4 axial directions. Particularly notable is the vector into the small pocket between His164, Met165 and Asp187, exploited by three of the fragments (Z1220452176 (x0104), Z219104216 (x0305) and Z509756472 (x1249)) with fluoro and cyano substituents (Figure 3).

Of the four fragments exploring subsite S3, three contain an aromatic ring with a sulfonamide group forming hydrogen bonds with Gln189 and pointing out of the active site towards the solvent interface (Figure 3). These hits have expansion vectors suitable for exploiting the same His164/Met165/Asp187 pocket mentioned above.

The experiment revealed one notable conformational variation, which was exploited by one fragment only (Z369936976 (x0397); Figure 4): a change in the sidechains of the key catalytic residues His41, Cys145 alters the size and shape of subsite S1′ and thus the link to subsite S1. This allows the fragment to bind, uniquely, to both S1 and S1′. In S1, the isoxazole nitrogen hydrogen-bonds to His163, an interaction that features in several other hits; and in S1′, the cyclopropyl group occupies the region sampled by the covalent fragments. Notably, the N-methyl group offers a vector to access the S2 and S3 subsites.

**Figure 4.**
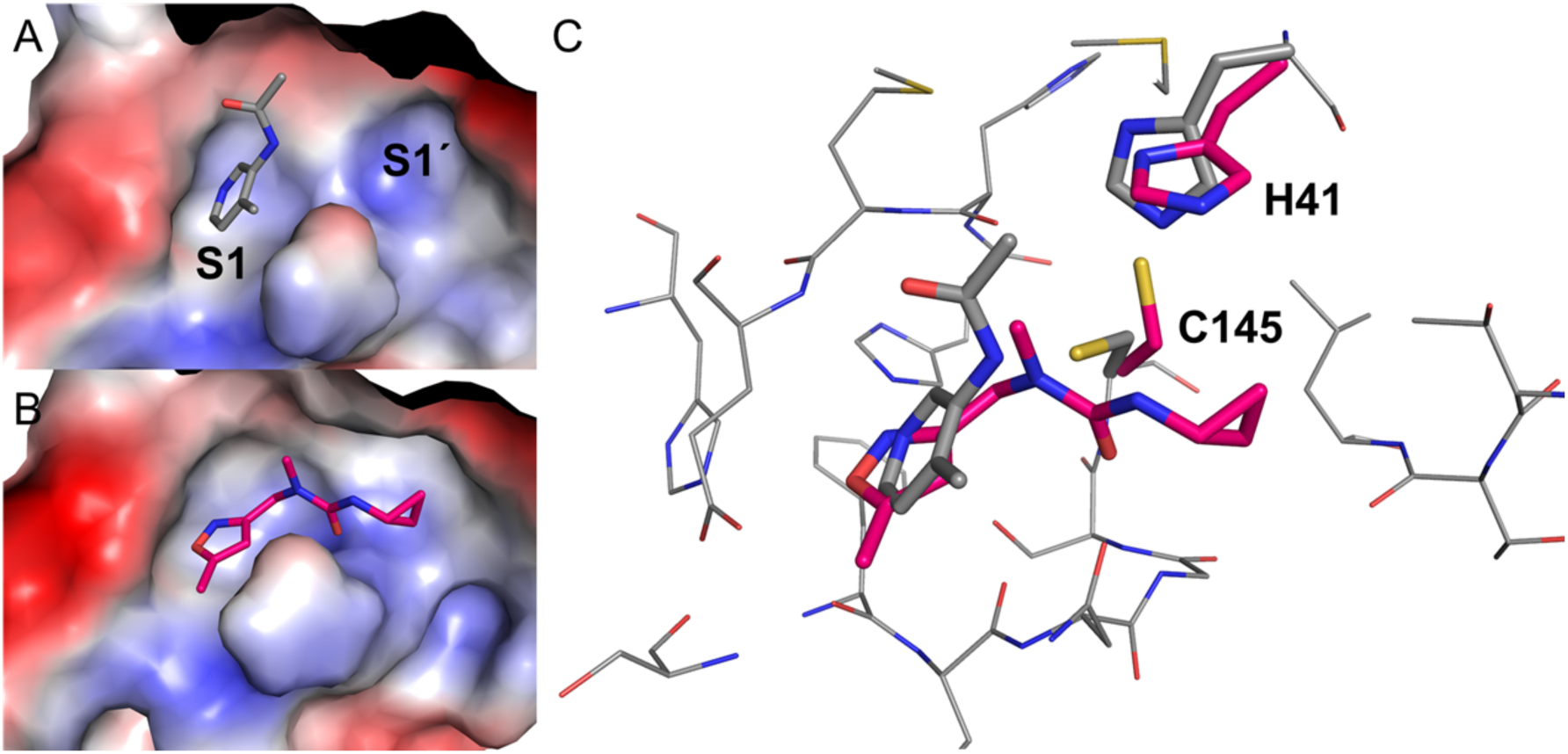
Plasticity of S1’ is revealed by fragment Z369936976 (x0397). Comparing the electrostatic surfaces of Z1129283193 (x0107) (A), the most commonly observed conformation, with that of Z369936976 (x0397) (B) shows how the shape of S1 and S1’ can change. C: Sidechain movement of catalytic residues Cys145 and His41 upon binding of Z369936976 (x0397, magenta) compared to Z1129283193 (x0197, grey).

#### Dimer interface fragments

It is established that the biological unit for similar viral proteases, such as the SARS-CoV-1 protease is a dimer (Chou et al., 2004), and that mutations at the dimer interface can disrupt proteases activity (Chen et al., 2008, Hsu et al., 2005) even at long range (Barrila et al., 2006). Thus, compounds that interfere with dimerization might serve as quasi-allosteric inhibitors of protease activity. In this study three compounds bound at the dimer interface.

Fragment Z1849009686 (x1086; Figure 5A) binds in a hydrophobic pocket formed by the sidechains of Met6, Phe8, Arg298 and Val303. It also mediates two hydrogen bonds to the sidechain of Gln127 and the backbone of Met6. Its binding site is less than 7 Å away from Ser139, whose mutation to alanine in SARS-CoV-1 protease reduced both dimerization and protease activity by about 50% (Chen et al., 2008, Hu et al., 2009). Z264347221 (x1187, Figure 5B) binds similarly in a hydrophobic pocket made by Met6, Phe8 and Arg298 in one of the protomers, extending across the dimer interface to interact with Ser123, Tyr118 and Leu141 of the second protomer, including hydrogen bonds with the sidechain and backbone of Ser123. Finally, POB0073 (x0887; York 3D library; Figure 5C), binds only 4 Å from Gly2 at the dimer interface and is encased between Lys137 and Val171 of one protomer and Gly2, Arg4, Phe3, Lys5 and Leu282 of the second, including two hydrogen bonds with the backbone of Phe3.

**Figure 5.**
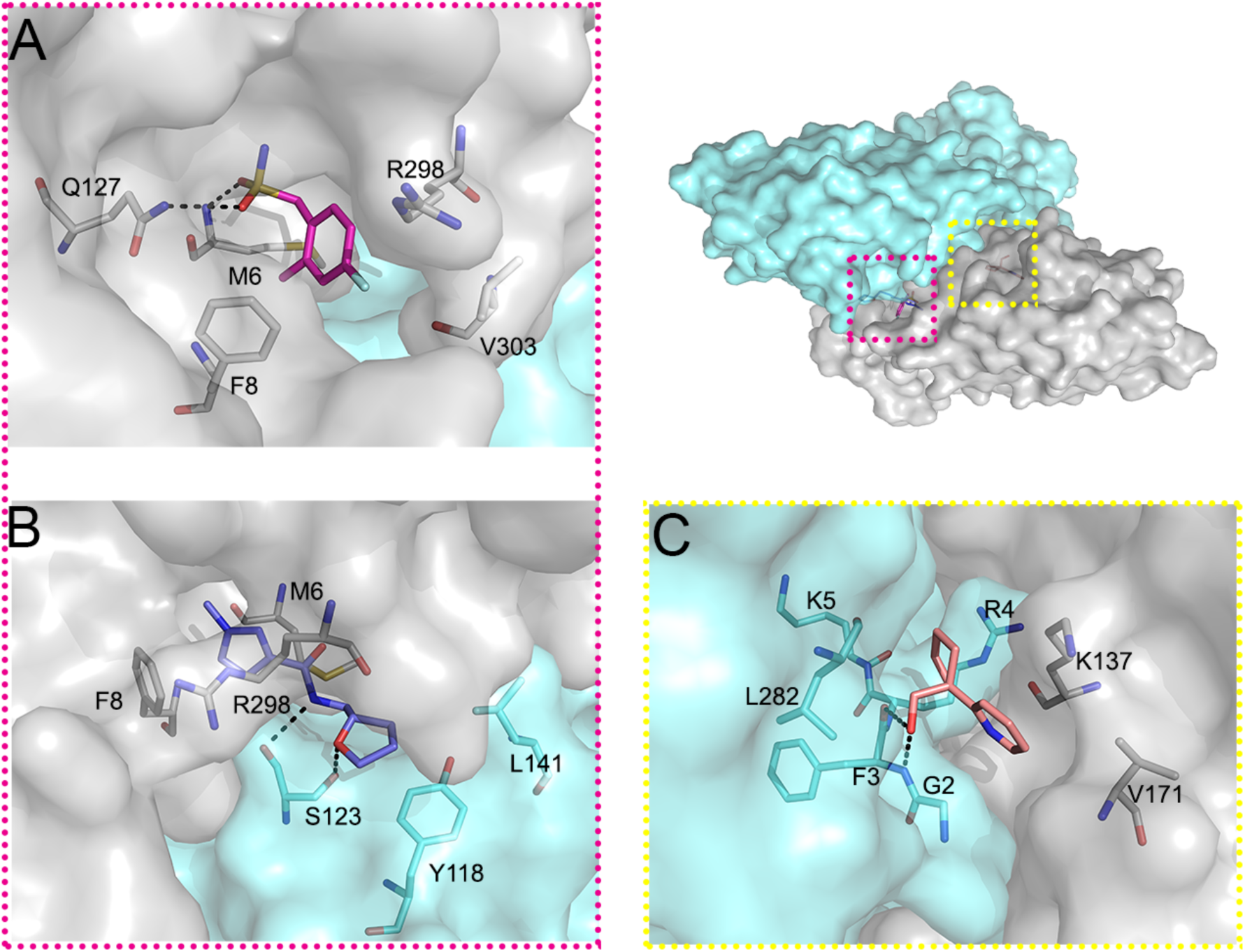
Fragments at dimer interface indicate opportunities for allosteric modulation. The overview shows the surface of the M^pro^ dimer, the protomers in grey and cyan. Fragments and surrounding residues are shown as sticks and hydrogen bonds in dashed black lines. A. Z1849009686 (x1086). B. Z264347221 (x1187). C. POB0073 (x0887).

### Covalent fragment hits reveal several tractable series

The screen further yielded 48 structures of fragments covalently bound to the nucleophilic active site Cys145, and sample subsite S1’. The majority (44) fall into series explored in the mass spectrometry experiment and the remainder came from other libraries.

#### Electrophile fragments

In all structures with bound electrophiles, the *N*-chloroacetyl carbonyl oxygen atom forms either two or three hydrogen bonds with the backbone amide hydrogens of Gly143, Ser144 or Cys145 (Figure 6 A-C). All three compounds containing the *N*-chloroacetyl piperidinyl-4-carboxamide motif (Figure 6A) adopt a similar binding mode pointing towards the S2 pocket, and one (PCM-0102389, x1358) is able to form an additional hydrogen bond with the side chain of Asn142.

**Figure 6.**
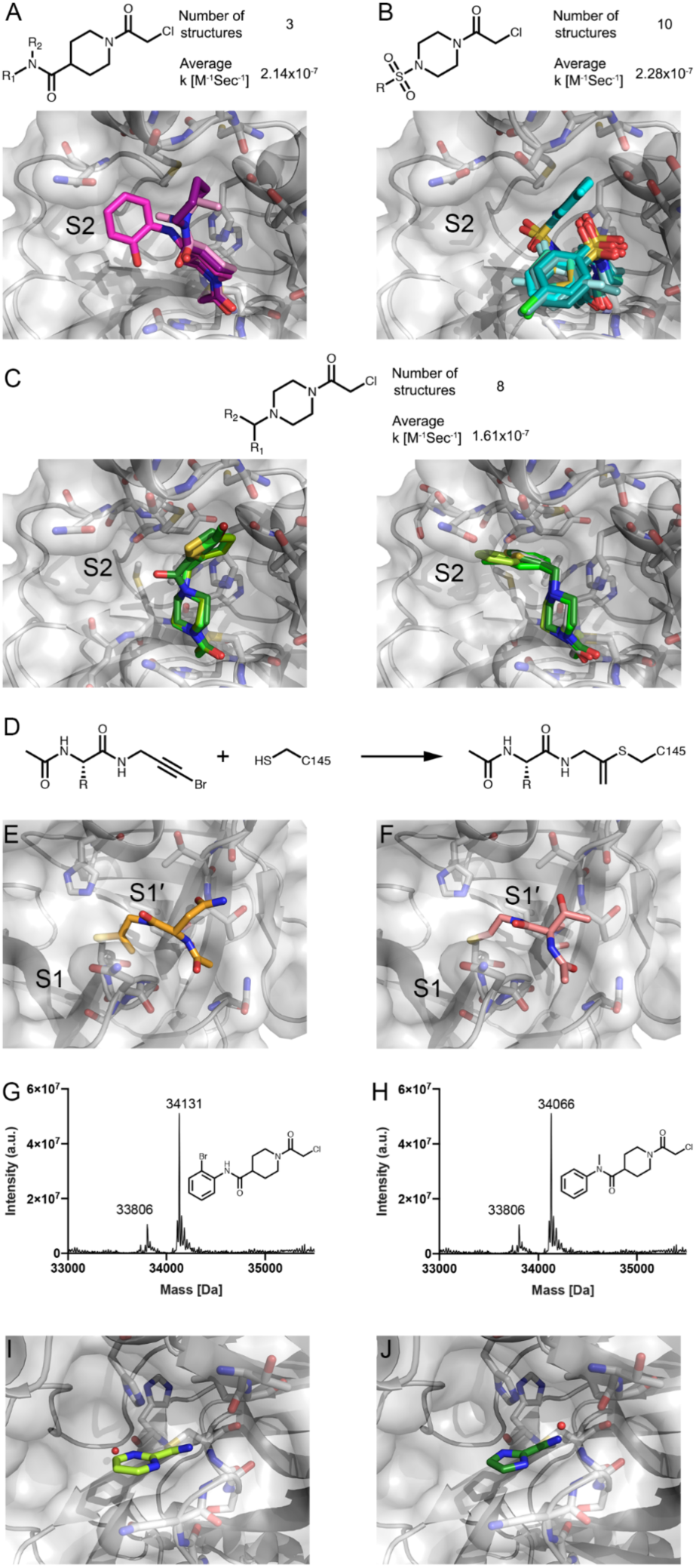
Covalent fragments are anchored at Cys145 and sample different regions of the orthosteric M^pro^ binding pocket. A. Fragments containing *N*-chloroacetyl piperidinyl-4-carboxamide motif. B. Fragments containing *N*-chloroacetyl-*N*’-sulfonamido-piperazine motif. C. Fragments containing *N*-chloroacetyl-*N*’-carboxamido- and *N*-chloroacetyl-*N*’-heterobenzyl-piperazine in two binding modes. D: Reaction schema of the unexpected covalent modification to Cys145 by PepLites hits. E. Threonine PepLite (NCL-00025058 (x0978)) bound covalently to active site cysteine. F: Asparagine PepLite (NCL-00025412 (x0981)) bound to active site cysteine. Labelling of M^pro^ by 2^nd^ generation compounds proven by intact protein LC-MS: G. labelling by PG-COV-35; H. labelling by PG-COV-34. Covalently bound cyclic electrophiles: I. Cov_HetLib 030 (x2097) and J. Cov_HetLib 053 (x2119).

Compounds with the *N*-chloroacetyl-*N*’-sulfonamido-piperazine motif (Figure 6B) adopt a bent shape, pointing towards the S2 pocket where appropriate space-filling substituents are attached to the phenyl moiety (PCM-0102353 (x1336) and PCM-0102395 (x0774)); otherwise, they point towards the solvent. Most of the latter 8 structures feature a halophenyl moiety which resides closely to Asn142, hinting at weak halogen-mediated interactions (Kuhn et al., 2019).

Eight compounds with a *N*-chloroacetyl-*N*’-carboxamido- and *N*-chloroacetyl-*N*’-heterobenzyl-piperazine motif crystallized in one binding mode with respect to the piperazinyl moiety (Figure 6C) (with one exception, PCM-0102287 (x0830)). Two structures (PCM-0102277 (x1334), PCM-0102169 (x1385)) with a 5-halothiophen-2-ylmethylene moiety exploit lipophilic parts of S2, which is also recapitulated by the thiophenyl moiety in an analogous carboxamide (PCM-0102306 (x1412)). The other five structures point mainly to S2, offering an accessible growth vector towards the nearby S3 pocket.

A series of compounds containing a N-chloroacetyl piperidinyl-4-carboxamide motif showed promising binding modes. To follow up on these compounds we performed a rapid second-generation compound synthesis. Derivatives of this chemotype were accessible in mg-scale by reaction of N-chloroacetyl piperidine-4-carbonyl chloride with various in-house amines, preferably carrying a chromophore to ease purification. These new compounds were tested by intact protein mass-spectrometry to assess protein labelling (5 uM compound; 1.5h incubation, RT; Table S1b). Amides derived from non-polar amines mostly outcompeted their polar counterparts, hinting at a targetable lipophilic sub-region in this direction. The two amides with the highest labelling PG-COV-35 and PG-COV-34 (figure 6G,H) highlight the potential for further synthetic derivatization by amide N-alkylation or cross-coupling, respectively.

#### PepLites

The screen revealed unexpected covalent warheads from the series of 3-bromoprop-2-yn-1-yl amides of N-acylamino acids. Colloquially termed PepLites (Noble and Waring), this library was developed to map non-covalent interactions of amino acid sidechains in protein-protein interaction hotspots, with the acetylene bromine intended, as for FragLites (Bauman et al., 2016, Wood et al., 2019), as detection tag by anomalous dispersion in X-ray crystallography. However, bromoalkynes can also act as covalent traps for activated cysteine thiols (Mons et al., 2019) (figure 6D).

Two PepLites, containing threonine (NCL-00025058 (x0978)) and asparagine (NCL00025412 (x0981)) bound covalently to the active site cysteine (Cys145), forming a thioenolether via C-2 addition with loss of bromine (Figure 6E,F). The covalent linkage was unexpected and evidently the result of significant non-covalent interactions, specific to these two PepLites, that position the electrophile group for nucleophilic attack. We note the side-chains make hydrogen-bonding interactions with various backbone NH and O atoms of Thr26 and Thr24; in the case of threonine, it was the minor 2R,3R diastereomer (corresponding to D-allothreonine) that bound. The only other PepLite observed (tyrosine, NCL-00024905 (x0967)) bound non-covalently to a different subsite.

The highlighted structure activity relationships is important for further optimisation. Bromoalkynes have intrinsic thiol reactivity that is lower than that of established acrylamide-based covalent inhibitors (Mons et al., 2019), which is in general desirable. The geometry of the alkyne and its binding mode also suggest that it could be replaced by reversible covalent groups such as nitriles, which would be guided by the same non-covalent interactions but are better established as cysteine protease inhibitors.

#### Heterocyclic electrophiles

Two covalent hits (2-cyano pyrimidine (Cov_HetLib 030 (x2097)) and 2cyano-imidazole (Cov_HetLib 053 (x2119) came from a library of small heterocyclic electrophiles (Keeley et al., 2019). These are essentially covalent MiniFrags (O’Reilly et al., 2019), comprising five and six-membered nitrogen containing heterocycles with electron-withdrawing character that activates small electrophilic substituents (halogenes, acetyl, vinyl and nitrile groups).

Both hits bound to Cys145 through an imine (Figure 6I,J), positioned by a local hydrogen bond network involving imine and heterocyclic N atoms. One of these free amines provides an immediate growth vector towards to catalytic pocket. The compounds have reasonable stability in water (Keeley et al., 2018) and limited reactivity against GSH (t1/2= 2.2 and 52.3 h, respectively), well above suggested reactivity limits (Fuller et al., 2016). They are also inactive against various covalent targets (HDAC8, MAO-A, MAO-B, MurA) and benchmark proteins.

## Discussion

The data presented herein provides many clear routes to developing potent inhibitors against SARS-CoV-2. The bound fragments comprehensively sample all subsites of the active site revealing diverse expansion vectors, and the electrophiles provide extensive, systematic as well as serendipitous, data for designing covalent compounds.

It is widely accepted that new small molecule drugs cannot be developed fast enough to help against COVID-19. Nevertheless, as the pandemic threatens to remain a long-term problem and vaccine candidates do not promise complete and lasting protection, antiviral molecules will remain an important line of defence. Such compounds will also be needed to fight future pandemics (Hilgenfeld, 2014). Our data will accelerate such efforts: therapeutically, through design of new molecules and to inform ongoing efforts at repurposing existing drugs; and for research, through development of probe molecules (Arrowsmith et al., 2015) to understand viral biology. One example is the observation that fragment Z1220452176 (x0104) is a close analogue of melatonin, although in this case, it is unlikely that melatonin mediates direct antiviral activity through inhibition of M^pro^, given its low molecular weight; nevertheless, melatonin is currently in clinical trials to assess its immune-regulatory effects on COVID19 (Clinicaltrials.gov identifier NCT04353128).

In line with the urgency, results were made available online immediately for download. Additionally, since exploring 3D data requires specialised tools (Ferla et al., 2020, Lee et al., 2011), hits were made accessible on the Fragalysis webtool (https://fragalysis.diamond.ac.uk) that allows easy exploration of the hits in interactive 3D.

We have previously demonstrated the benefits of merging covalent and non-covalent fragments to make dramatic improvements in potency (Resnick et al., 2019). Our dataset offers numerous opportunities and some conservative examples are shown in figure 7. These can be expected to result in potent M^pro^ binders and compound synthesis is ongoing.

**Figure 7.**
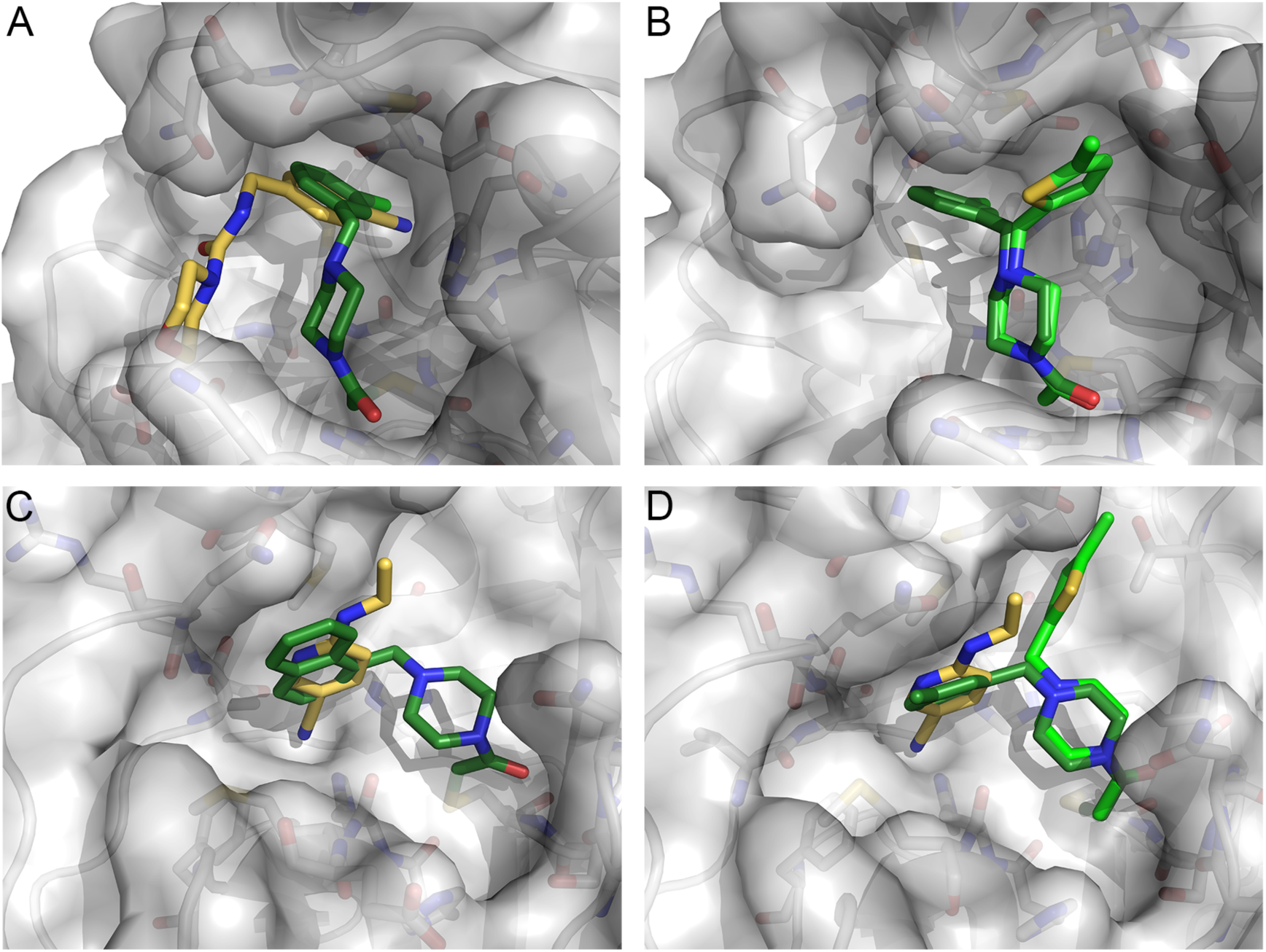
Fragment merging opportunities can be directly inferred from many hits. Covalently-bound fragments are in green shades, and non-covalent fragments in yellow A. Overlay of Z509756472/x1249 and PCM-0102269/x0770. B. Overlay of PCM-0102277/x1334 and PCM-0102269/x0770. C. Overlay of PCM-0102287/x0830 and Z219104216/x0305. D. Overlay of PCM-0102340/x0692, PCM-0102277/x1334 and Z219104216/x0305.

Collectively, the covalent hits provide rational routes to inhibitors of low reactivity and high selectivity. Rationally designed covalent drugs are gaining traction, with many recent FDA approvals (Singh et al., 2011, Bauer, 2015). Their design is based on very potent reversible binding, that allows precise orientation of a low reactivity electrophile, so that formation of the covalent bond is reliant on binding site specificity, with minimal off-targets. (De Cesco et al., 2017, Zhang et al., 2019, Baillie, 2016). For this reason, covalent inhibitors are expunged from high-throughput screening libraries and are typically considered as PAINS compounds (Sirois et al., 2005, Baell and Nissink, 2018, Baell and Holloway, 2010). The challenge of tuning reactivity, and the danger of reactivity-based artefacts, are likely to be particularly marked for the highly reactive nucleophiles such as the catalytic cysteine of many proteases. This is evidenced by the very high hit-rate we saw in our preliminary electrophiles screen in which more than 150 fragments labelled M^pro^ by >50%. Robust characterization of the fragments’ reactivity (Resnick et al., 2019), and continuous evaluation of general thiol reactivity in the selection of lead series and during hit-to-lead optimization can address this challenge.

The scale of this experiment, particularly the diversity of libraries and density of results, likely sets a new benchmark for ensuring a crystal-based fragment screen accelerates progression of hits. Even cursory inspection of the fragment structures indicates a very large “merge space”, i.e. the collection of compounds that can be designed directly from spatial juxtapositions of fragments. Such merges, which can be made to populate all four subsites, might achieve potency synergistically, because the observed interactions can be assumed to be in near-optimal configurations, given how few there are per fragment. A thorough exploration of merge space might be best achieved formulaically, using computational workflows that additionally filter undesirable molecular properties, assess synthetic tractability and predict binding affinity. However, such integrated approaches are not currently available in the public domain. We hope this dataset will help spur their development and testing.

Another promising effort to explore the potential of this premise is the COVID Moonshot project (https://covid.postera.ai/covid), where a selection of merges will be experimentally tested, with data promptly made public. We trust that this resource will enable the development of many new tools, approaches and ultimately viable treatment candidates for COVID19.

## Materials and Methods

### Protein Expression

Multiple transformant colonies were used to inoculate a starter culture supplemented with 100 µg/ml Carbenicillin. The culture was then grown to log phase for approximately 8 h. 10 ml of the starter culture was used to inoculate 1 litre of Auto Induction Medium supplemented with 10 ml of glycerol and 100 µg/ml Carbenicillin. The cultures were grown at 37 °C, 200 rpm for 5 h then switched to 18 °C, 200 rpm for 10 h. The cells were harvested by centrifugation and stored at −80 °C

### Protein purification

Cells were resuspended in 50 mM Tris pH 8, 300 mM NaCl, 10 mM Imidazole, 0.03 μg/ml Benzonase. The cells were disrupted on a high-pressure homogeniser (3 passes, 30 kpsi, 4 °C). The lysate was clarified by centrifugation at 50,000 g. The supernatant was then applied to a Nickel-NTA gravity column and washed and eluted with 50 mM Tris pH 8, 300 mM NaCl, and 25-500 mM imidazole pH 8. N-terminal His tagged HRV 3C Protease was then added to the eluted protein at 1:10 w/w ratio. The mixture was then dialysed overnight at 4 °C against 50 mM Tris pH 8, 300 mM NaCl, 1 mM TCEP. The following day, the HRV 3C protease and other impurities were removed from the cleaved target protein by reverse Nickel-NTA. The relevant fractions were concentrated and applied to an S200 16/60 gel filtration column equilibrated in 20 mM Hepes pH 7.5, 50 mM NaCl buffer. The protein was concentrated to 30 mg/ml using a 10 kDa MWCO centrifugal filter device.

### Crystallisation and structure determination

Protein was thawed and diluted to 5 mg/ml using 20 mM Hepes pH 7.5, 50 mM NaCl. The sample was centrifuged at 100 000 g for 15 minutes. Initial hits were found in well F2 of the Proplex crystallisation screen, 0.2 M LiCl, 0.1M Tris pH 8, 20% PEG 8K. These crystals were used to prepare a seed stock by crushing the proteins with a pipette tip, suspending in reservoir solution and vortexing for 60 s in the reservoir solution with approximately 10 glass beads (1.0mm diameter, BioSpec products). Adding DMSO to the protein solution to a concentration of 5% and performing microseed matrix screening, many new crystallisation hits were discovered in commercial crystallisation screens. Following optimisation, the final crystallisation condition was 11% PEG 4K, 6% DMSO, 0.1M MES pH 6.7 with a seed stock dilution of 1/640. The seeds were prepared from crystals grown in the final crystallisation condition. The drop ratios were 0.15 µl protein, 0.3 µl reservoir solution, 0.05 µl seed stock. Crystals were grown using the sitting drop vapor diffusion method at 20 °C and appeared within 24 hours.

Initial diffraction data was collected on beamline I04 at Diamond Light Source on a crystal grown in 0.1 M MES pH 6.5, 5% PEG6K, cryoprotected using 30% PEG400. Data were processed using Dials (Winter et al., 2018) via Xia2 (Winter et al., 2013). The dataset was phased with the SARS-CoV-2 M^pro^ in complex with the N9 inhibitor crystal structure (PDB:6LU7) using Molrep in CCP4i2. Further datasets were collected on I04-1 at Diamond Light Source on crystals grown using the 0.1 M MES pH 6.5, 15% PEG4K, 5% DMSO condition. To create a high-resolution dataset, datasets from 7 crystals were scaled and merged using Aimless (Evans and Murshudov, 2013). Crystal structures were manually rebuilt in Coot (Emsley et al., 2010) and refined using Refmac (Murshudov et al., 2011) and Buster (Bricogne et al., 2017). This structure is deposited in the PDB under ID 6YB7.

### Electrophile fragment LC/MS screen

2 µM M^pro^ was incubated in 50 mM Tris pH 8 300 mM NaCl for 1.5 hours at 25 °C. For initial electrophile fragment library screen, 30 µl protein with pools of 4-5 electrophile fragments, 7.5 nL each from 20 mM DMSO stocks and for other runs 50 µl protein with 0.5 µl compounds from 0.5 mM DMSO stocks. The reaction was quenched by adding formic acid to 0.4% final concentration. The LC/MS runs were performed on a Waters ACUITY UPLC class H instrument, in positive ion mode using electrospray ionization. UPLC separation used a C4 column (300 Å, 1.7 μm, 21 mm × 100 mm). The column was held at 40 °C and the autosampler at 10 °C. Mobile solution A was 0.1% formic acid in water, and mobile phase B was 0.1% formic acid in acetonitrile. The run flow was 0.4 mL/min with gradient 20% B for 4 min, increasing linearly to 60% B for 2 min, holding at 60% B for 0.5 min, changing to 0% B in 0.5 min, and holding at 0% for 1 min. The mass data were collected on a Waters SQD2 detector with an m/z range of 2−3071.98 at a range of 1000−2000 m/z. Raw data were processed using openLYNX and deconvoluted using MaxEnt. Labelling assignment was performed as previously described (Resnick et al., 2019).

### Fragment Screening

Fragments were soaked into crystals as previously described (Collins et al., 2017), by adding dissolved compound directly to the crystallisation drops using an ECHO liquid handler (final concentration 10% DMSO); drops were incubated for approximately 1 hour prior to mounting and flash freezing in liquid nitrogen. The following libraries were screened: the DSi-poised library (Enamine), a version of the poised library (Cox et al., 2016); a version of the MiniFrags library (O’Reilly et al., 2019) assembled in-house; the FragLites library (Wood et al., 2019); a library of shape-diverse 3D fragments (“York3D”) (Downes et al., 2020); heterocyclic electrophiles (Keeley et al., 2019); and the SpotFinder library (Bajusz and Keserü). All fragments were in 100% DMSO at varying stock concentrations, detailed at https://www.diamond.ac.uk/Instruments/Mx/Fragment-Screening/Fragment-Libraries.html).

Electrophile fragments identified by mass spectrometry were soaked by the same procedure as the other libraries, but in addition, they were also co-crystallised in the same crystallisation condition as for the apo structure. The protein was incubated with 10 to 20-fold excess compound (molar ratio) for approximately 1h prior to the addition of the seeds and reservoir solution (following Resnick *et al* (Resnick et al., 2019)).

Data were collected at the beamline I04-1 at 100K and processed to a resolution of approximately 1.8 Å using XDS (Kabsch, 2010) and either xia2 (Winter et al., 2013), autoPROC (Vonrhein et al., 2011) or DIALS (Winter et al., 2018). Further analysis was performed with XChemExplorer (Krojer et al., 2017): electron density maps were generated with Dimple (Keegan et al., 2015); ligand-binding events were identified using PanDDA (Pearce et al., 2017) (both the released version 0.2 and the pre-release development version (https://github.com/ConorFWild/pandda)); ligands were modelled into PanDDA-calculated event maps using Coot (Emsley et al., 2010); restraints were calculated with ACEDRG or GRADE (Long et al., 2017, Smart et al., 2010); and structures were refined with Refmac (Murshudov et al., 2011) and Buster (Bricogne et al., 2017). A more thorough description of the PanDDA analysis is provided in the supplementary information.

Coordinates, structure factors and PanDDA event maps for all data sets are deposited in the Protein Data Bank under group deposition ID G_1002135, G_1002151, G_1002152, G_1002153, G_1002156 and G_1002157. Data collection and refinement statistics are summarised in supplementary table 4. The ground-state structure and all corresponding datasets are deposited under PDB ID 5R8T.

### Synthesis of *N*-chloroacetyl-piperidine-4-carboxamides

*N*-chloroacetyl piperidine-4-carbonyl chloride was prepared as a stock solution in dry DCM under an atmosphere of N_2_. Briefly, deprotecting *N*-Boc isonepecotic acid in 50% TFA in DCM (v/v) at RT for 2 h yielded the corresponding TFA salt after evaporation of all volatiles. The crude TFA salt was then re-dissolved in DCM, treated with Et_3_N (2 equiv.), followed by the addition of chloroacetic anhydride (1 equiv.). The reaction mixture was stirred overnight at RT, washed with water, the organic phase dried over MgSO_4_, filtered, and all volatiles removed by rotary evaporation. The crude N-chloroacetyl piperidine-4-carboxylic acid was refluxed in excess neat SOCl_2_ (gas evolution and a colour change to red occurs) for 1 h, followed by removal of excess SOCl_2_ in vacuum into a liquid nitrogen-cooled trap. The remaining residue was dried by rotary evaporation, placed under an atmosphere of nitrogen and dissolved in dry DCM to give a stock solution of approx. 0.489 M (based on theoretical yield over three steps), which was immediately used.

The corresponding amides were prepared by addition of the acid chloride (1 equiv.) as a DCM solution to the pertinent amines (1 equiv.) in presence of pyridine (1 equiv.) in DCM. Heterogeneous reaction mixtures were treated with a minimal amount of dry DMF to achieve full solubility. After stirring the reaction mixtures overnight, the solvents were removed in by rotary evaporation, re-dissolved in 50% aq. MeCN (and a minimal amount of DMSO to achieve higher solubility), followed by purification by (semi-)preparative RP-HPLC in mass-directed automatic mode or manually.

### Synthesis of PepLites

HATU (1.5 eq.), DIPEA (3.0 eq.) and the acid starting material (1.5 eq.) were dissolved in DMF (3 - 6 mL) and stirred together at room temperature for 10 min. 3-Bromoprop-2-yn-1-amine hydrochloride was added and the reaction mixture was stirred at 40 °C overnight. The reaction mixture was allowed to cool to room temperature, diluted with EtOAc or DCM and washed with saturated aqueous sodium bicarbonate solution, brine and water. The organic layer was dried over MgSO_4_, filtered and evaporated to afford crude product. The crude product was then purified by either normal or reverse phase chromatography.

### *tert*-Butyl (3-bromoprop-2-yn-1-yl)carbamate

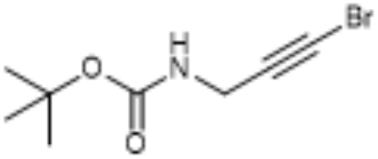

A solution of KOH (2.7 g, 48 mmol) in water (15 mL) was added dropwise to a solution of *N*-bocpropargylamine (3.0 g, 19 mmol) in MeOH (45 mL) at 0 °C under nitrogen. The resulting solution was stirred at 0 °C for 10 min then bromine (1.1 mL, 21 mmol) was added dropwise. The reaction mixture was allowed to warm to room temperature and was stirred at room temperature for 24 h. The reaction mixture was diluted with water and extracted with diethyl ether. The organic extracts were combined, dried over MgSO_4_ and evaporated to afford crude product. The crude product was purified by flash silica chromatography, elution gradient 0 – 10% EtOAc in petroleum ether. Pure fractions were evaporated to dryness to afford *tert*-Butyl (3-bromoprop-2-yn-1-yl)carbamate (3.5 g, 79%) as a white solid. *R*_f_ = 0.34 (10% EtOAC in petroleum ether); m.p. 108 - 110 °C; IR ν_max_ (cm^-1^) 3345, 2982, 2219, 2121, 2082; ^1^H NMR (500 MHz, DMSO-*d*6) δ 1.39 (s), 3.76 (d, *J* = 5.9 Hz), 7.30 (d, *J* = 6.1 Hz). LCMS m/z ES^+^ [M-Boc+H]^+^ 133.9.

### 3-Bromoprop-2-yn-1-amine hydrochloride

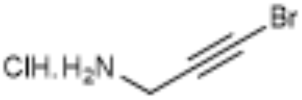

*tert*-Butyl (3-bromoprop-2-yn-1-yl)carbamate (1.1 g, 4.7 mmol) was dissolved in 4M HCl in dioxane (30 mL). The reaction mixture was stirred at room temperature for 2 h then evaporated to dryness to afford 3-bromoprop-2-yn-1-amine hydrochloride (0.79 g, 99%) as a yellow solid. m.p. 169 °C; IR ν_max_ (cm^-1^) 2856, 2629, 2226, 2121, 2074; ^1^H NMR (500 MHz, DMSO-*d*_6_) δ 3.78 (s, 2H), 8.48 (s, 3H); LCMS m/z ES^+^ [M+H]^+^ 171.9.

### (2*S*,3*R*)-2-Acetamido-*N*-(3-bromoprop-2-yn-1-yl)-3-(*tert*-butoxy)butanamide

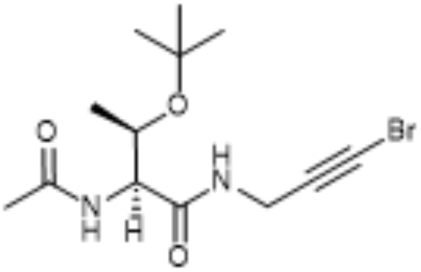

(2S,3S)-2-Acetamido-N-(3-bromoprop-2-yn-1-yl)-3-(tert-butoxy)butanamide was synthesized according to General procedure A using (2*S*,3*R*)-2-acetamido-3-(*tert*-butoxu)butanoic acid (0.41 g, 1.9 mmol). The crude product was purified by flash silica chromatography, elution gradient 0 – 10% MeOH in DCM. Pure fractions were evaporated to dryness to afford (2*S*,3*S*)-2-acetamido-*N*-(3-bromoprop-2-yn-1-yl)-3-(*tert*-butoxy)butanamide (0.20 g, 42%) as a white solid. R_f_ = 0.46 (10% MeOH in DCM); mp: 180 – 183 °C; IR ν_max_ (cm^-1^) 3271, 3078, 2969, 2935, 2222, 2113; ^1^H NMR (500 MHz, Methanol-*d*_4_) δ 1.16 (d, *J* = 6.2, 5.0 Hz), 1.21 (s, *J* = 3.9 Hz, 9H), 2.01 (s, 3H), 3.91 – 4.09 (m, 3H), 4.32 (d, *J* = 7.5 Hz, 1H); ^13^C NMR (126 MHz, Methanol-*d*_4_) δ 18.61, 21.15, 27.27, 28.90, 41.92, 58.81, 67.21, 74.16, 75.57, 171.19, 171.92; LCMS m/z ES+ [M+H]+ 333.2; calcd for C_13_H_21_^79^BrN_2_O_3_ 333.2260 [M(Br)+H]^+^ found 333.0808.

### (2*S*,3*R*)-2-Acetamido-*N*-(3-bromoprop-2-yn-1-yl)-3-hydroxybutanamide (threonine PepLite)

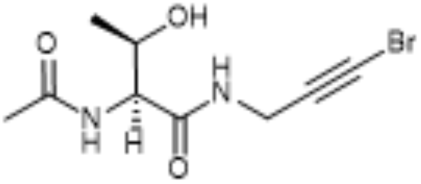

(2*S*,3*S*)-2-Acetamido-*N*-(3-bromoprop-2-yn-1-yl)-3-(*tert*-butoxy)butanamide (80 mg, 0.24 mmol) was dissolved in anhydrous DCM (20 mL) and TFA (10 mL) and 0 °C under nitrogen. The reaction mixture was allowed to warm to room temperature and was stirred at room temperature for 3 h then evaporated to dryness to afford crude product. The crude product was purified by flash silica chromatography, elution gradient 0 – 15% MeOH in DCM. Pure fractions were evaporated to dryness to afford (2*S*,3*S*)-2-acetamido-*N*-(3-bromoprop-2-yn-1-yl)-3-hydroxybutanamide (38 mg, 57%, 93% de) as a white solid. R_f_ = 0.34 (10% MeOH in DCM); mp: 189 – 192 °C; IR ν_max_ (cm^-1^) 3280, 3085, 2973, 2924, 2225, 2115; ^1^H NMR (500 MHz, Methanol-*d*_4_) δ 1.21 (d, *J* = 6.4 Hz, 3H), 2.03 (s, 3H), 3.97 – 4.06 (m, 3H), 4.33 (d, *J* = 6.5 Hz, 1H); ^13^C NMR (126 MHz, Methanol-*d*_4_) δ 18.21, 21.13, 29.00, 41.79, 58.69, 67.11, 75.41, 170.88, 172.00; LCMS m/z ES^+^ [M+H]^+^ 277.1; calcd for C_9_H_13_^79^BrN_2_O_3_ 277.1180 [M(Br)+H]^+^ found 277.0182.

### (*S*)-2-Acetamido-*N*^1^-(3-bromoprop-2-yn-1-yl)succinimide (asparagine PepLite)

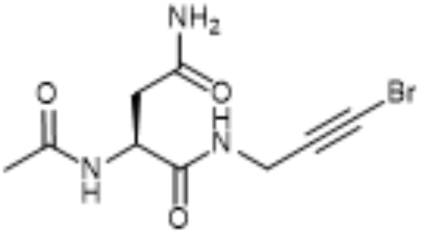

(*S*)-2-Acetamido-*N*^1^-(3-bromoprop-2-yn-1-yl)succinamide was synthesized according to General procedure A using (*s*)-2-acetamido-5-amino-5-oxobutanoic acid (155 mg, 0.89 mmol) and evaporating the reaction mixture to afford the crude product without aqueous work-up. The crude product was purified by flash silica chromatography, elution gradients 0 – 10% MeOH in DCM. Pure fractions were evaporated to dryness to afford (*S*)-2-acetamido-*N*^1^-(3-bromoprop-2-yn-1-yl)succinamide (50 mg, 30%) as a white solid. R_f_ = 0.18 (10% MeOH in DCM); mp: 173 °C (decomp); IR ν_max_ (cm^-1^) 3421, 3277, 3208, 3072, 2922, 2226, 2116; 1H NMR (500 MHz, Methanol-d4) δ 1.99 (s, 3H), 2.58 – 2.75 (m, 2H), 3.98 (d, J = 1.4 Hz, 2H), 4.71 (dd, J = 7.6, 5.7 Hz, 1H); 13C NMR (126 MHz, Methanol-d4) δ 22.57, 30.61, 37.83, 43.13, 51.54, 76.84, 173.04, 173.28, 174.81; LCMS m/z ES^+^ [M+H]^+^ 290.2.

## Acknowledgements

We would like to thank all the staff of Diamond Light Source for providing support and encouragement which allowed us to carry out this work during the COVID-19 lockdown. We’d also like to thank the Diamond MX group for their support and expertise, in particular David Aragão, Ralf Flaig, Dave Hall, Katherine McAuley and Mark Williams. We are grateful to AstraZeneca, Astex Pharmaceuticals, Lilly, Pfizer and Vernalis, University of York (TDD, SPJ) and the EU (Horizon 2020 program, Marie Skłodowska-Curie grant agreement No. 675899, FRAGNET) (AK and HFK) for support. The SGC is a registered charity (number 1097737) that receives funds from AbbVie, Bayer Pharma AG, Boehringer Ingelheim, Canada Foundation for Innovation, Eshelman Institute for Innovation, Genome Canada, Innovative Medicines Initiative (EU/EFPIA) [ULTRA-DD grant no. 115766], Janssen, Merck KGaA Darmstadt Germany, MSD, Novartis Pharma AG, Ontario Ministry of Economic Development and Innovation, Pfizer, São Paulo Research Foundation-FAPESP, Takeda, and Wellcome [106169/ZZ14/Z]. N.L. is the incumbent of the Alan and Laraine Fischer Career Development Chair. N.L. would like to acknowledge funding from the Israel Science Foundation (grant no. 2462/19), The Israel Cancer Research Fund, the Israeli Ministry of Science Technology (grant no. 3-14763), and the Moross Integrated Cancer Center. N.L. is also supported by the Helen and Martin Kimmel Center for Molecular Design, Joel and Mady Dukler Fund for Cancer Research, the Estate of Emile Mimran and Virgin JustGiving, and the George Schwartzman Fund.

