## Supplemental tables and figures for "Crystallographic and electrophilic fragment screening of the SARS-CoV-2 main protease"

### Supplementary Information

#### 1. PanDDA algorithm facilitates identification of weakly bound ligands

All datasets were analysed using the Pan Dataset Density Analysis (PanDDA) method(Pearce et al., 2017a). The PanDDA algorithm takes advantage of the large number of datasets collected during a fragment campaign to detect partial-occupancy ligands that are not visible in normal crystallographic maps (Fig. S1). PanDDA uses a statistical analysis to identify bound ligands, and then generates an “event map” for the bound state of the crystal. An event map approximates what would be observed if the ligand was bound at full occupancy and is generated by subtracting the unbound fraction of the crystal from the partial-occupancy dataset. While most of the fragments are also obvious in conventional maps, several ligands may appear unjustifiably modelled when only the corresponding 2mFo–DFc is inspected. Therefore, PanDDA event maps are included in the MMCIF file that can be downloaded from the PDB website. Additionally, each deposition of models analysed with PanDDA is accompanied by a separate deposition of the ground-state model which also contains structure factors of all the collected datasets and which can be used to reproduce the analysis. Finally, PanDDA models are refined as composite models consisting of the ligand-bound and confounding ground-state(Pearce et al., 2017b). But since these models can become rather complex and difficult to interpret, we deposited the bound-state only into the PDB. This might lead to slightly elevated R/Rfree values in some cases.

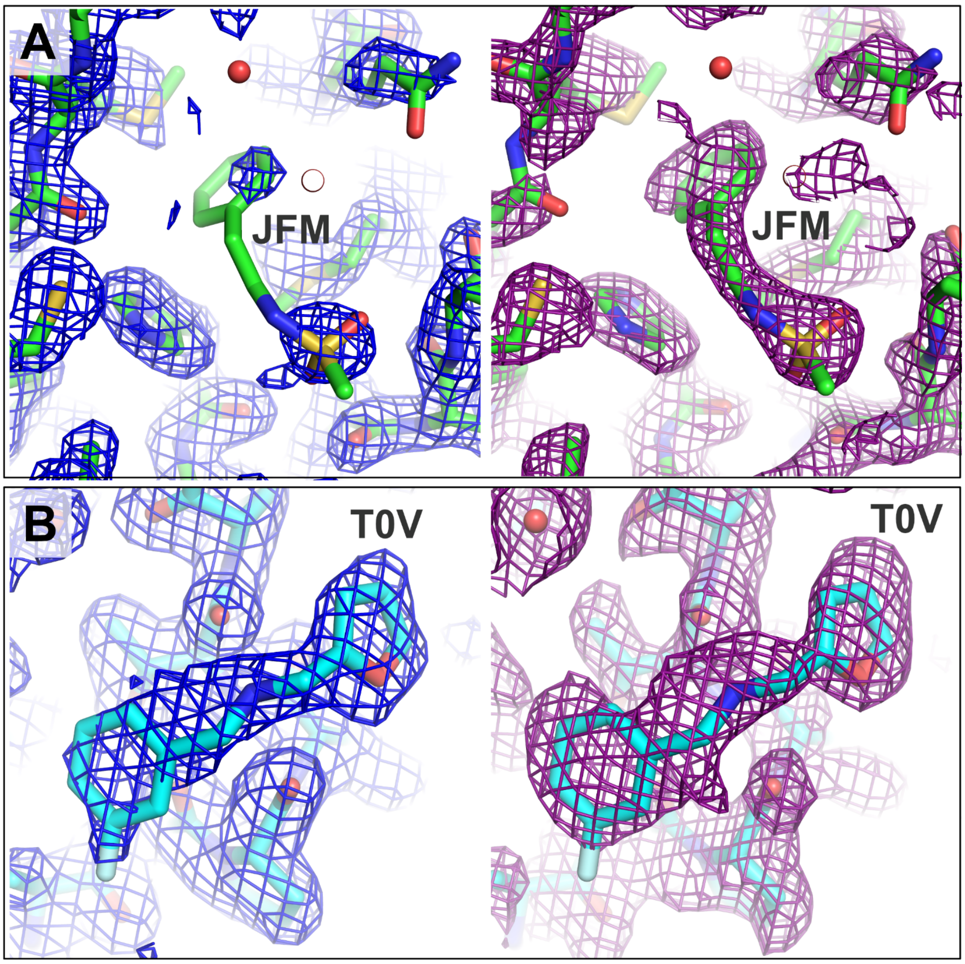


**Fig. S1. PanDDA event maps enable discovery of weakly bound fragments.** The figure shows two examples of a weakly and a strongly bound ligand. Conventional 2mFo–DFc maps contoured at 1 σ are shown on the left side (blue), whereas the corresponding PanDDA event map is shown on the right side (purple). (A) Example of a weakly bound ligand (JFM) and how the PanDDA event map served as evidence for the deposited structure (PDB ID 5R7Y). (B) Example of a ligand (T0V) that is clearly visible in a conventional crystallographic map (PDB ID 5RE8)

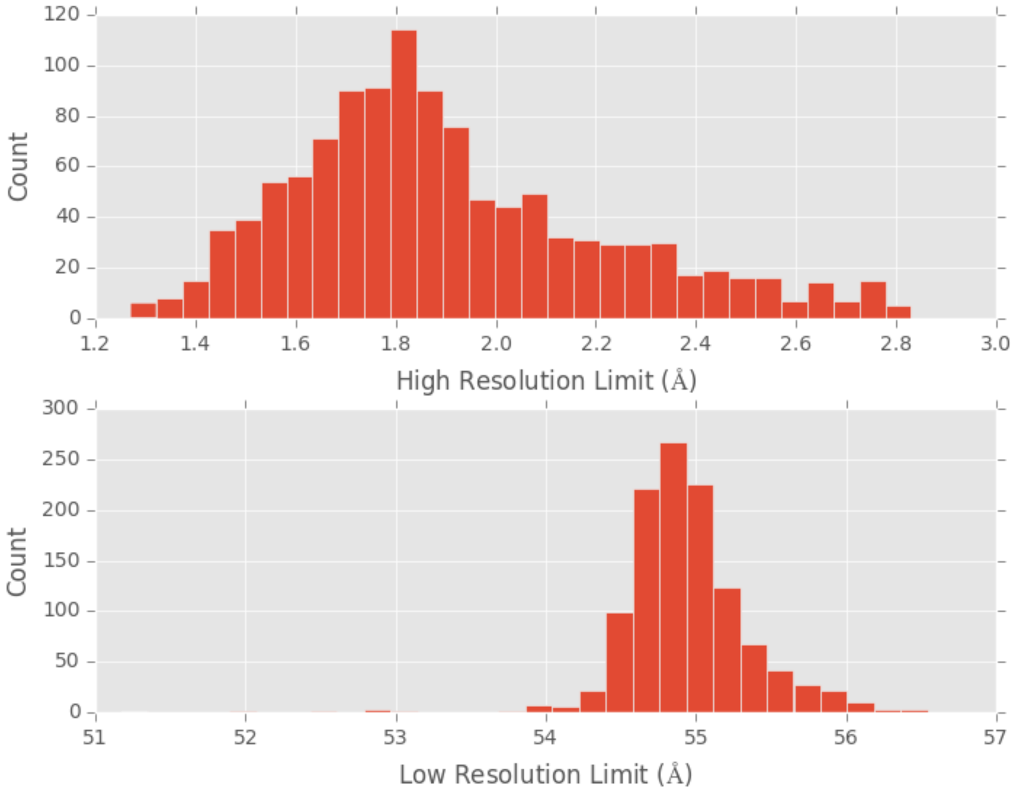


**Fig. S2. Distribution of resolutions of datasets analysed using PanDDA.**

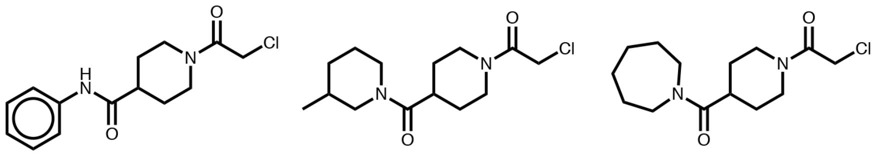

| | PCM-0102219 | PCM-0102933 | PCM-0102781 |
| --- | --- | --- | --- |
| Labeling | 85% | 44% | 37% |
| k [$M^{-1}Sec^{-1}$] | $2.80 \times 10^{-7}$ | $2.20 \times 10^{-7}$ | $1.86 \times 10^{-7}$ |

**Fig. S3. Labeling of compounds containing a *N*-chloroacetyl-*N*'-sulfonamido-piperazine or *N*-chloroacetylaniline motifs from electrophile fragment screen.**

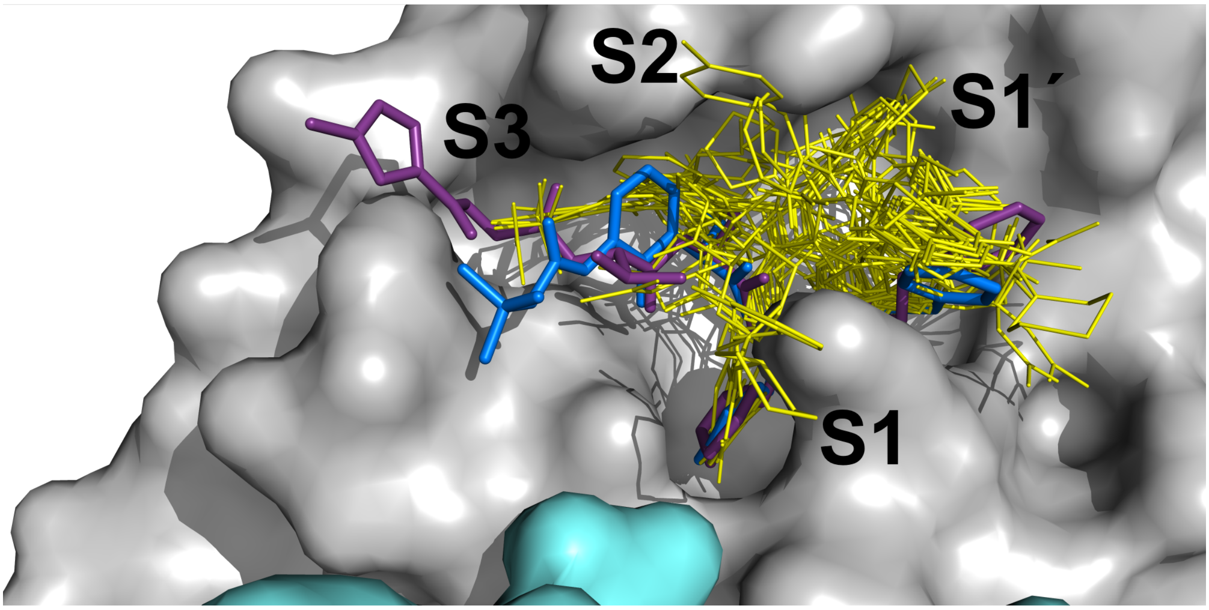


**Fig. S4. Surface representation of M$^{pro}$ dimer with fragment hits shown as yellow wires and peptide-based inhibitors from 6LU7 (purple) and 6Y2F (blue) shown as sticks.**

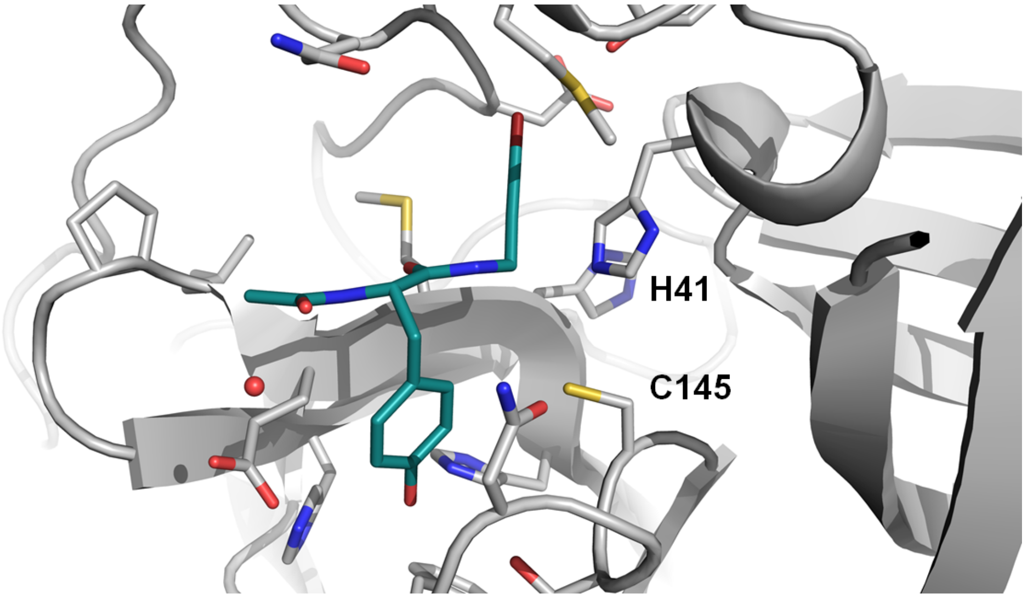


**Fig. S5. Tyrosine PepLite bound non-covalently in active site.**

**Table S1a. Mass spectrometry fragment screening.** 2 µM of Mpro was incubated with a pool of 5 electrophile fragments, 5 µM each at 25°C. After 1.5h the incubation was quenched by adding formic acid at a final concentration of 0.4%

EXCEL FILE TableS1a-b.xlsx

**Table S1b. Mass spectrometry fragment screening.** 2 µM of Mpro was incubated with 5 µM compound at 25 °C. After 1.5h the incubation was quenched by adding formic acid at a final concentration of 0.4%.

EXCEL FILE TableS1a-b.xlsx

**Table S2. Overview of all deposited structures from XChem fragment screening.** The table summarises all deposited $M^{pro}$-fragment structures and how internal crystal IDs map to compound IDs and respective fragment libraries. Furthermore, fragment binding locations are indicated: A – active site, B – active site covalent, C – dimer interface, D – surface, X – crystal contact; residues close to the binding site are shown in parenthesis. The table also provides a subjective assessment of the confidence the crystallographer has in the reliability of the model.

EXCEL FILE TableS2.xlsx

**Table S3. Summary of all screened fragments libraries.** The table gives an overview of which fragment libraries were screened, how many unique compounds each library contained and finally the number of hits that are currently deposited in the Protein Data Bank.

| Fragment Library | Source | Total number of compounds screened | Total number of compounds screened (unique) | Number of successful crystal mounts (unique) | Number of successful data collection (unique) | Number of PDB depositions |
|---|---|---|---|---|---|---|
| DSI poised | Diamond Light Source (UK) | 970 | 695 | 897 | 816 | 39 |
| Fraglites & Peplites | University of Newcastle (UK) | 53 | 53 | 53 | 42 | 4 |
| York3D | University of York (UK) | 106 | 106 | 102 | 91 | 2 |
| MiniFrags | Astex Therapeutics | 80 | 80 | 60 | 41 | 4 |
| electrophile cysteine covalent | Weizmann Institute of Science (Israel) | 1084 | 101 | 420 | 381 | 44 |
| SpotFinder | Hungarian Academy of Sciences | 114 | 96 | 112 | 107 | 1 |
| Heterocyclic electrophilic fragment library | Hungarian Academy of Sciences | 474 | 146 | 233 | 160 | 2 |

**Table S4. Crystallographic data collection and refinement statistics**

EXCE File TableS4.xlsx
